# Omicron-Specific and Bivalent Omicron-Containing Vaccine Candidates Elicit Potent Virus Neutralisation in the Animal Model

**DOI:** 10.1101/2022.02.11.480131

**Authors:** Asghar Abdoli, Hamidreza Jamshidi, Mohammad Taqavian, Mehdi Lari Baghal, Hasan Jalili

## Abstract

**Background:** Omicron variant (B. 1.1.529) is able to escape from naturally acquired and vaccine-induced immunity, which mandates updating the current COVID-19 vaccines. Here, we investigated and compared the neutralising antibody induction of the ancestral variant-based BIV1-CovIran vaccine, the Omicron variant-based BIV1-CovIran Plus vaccine, and the novel bivalent vaccine candidate, BBIV1-CovIran, against the Omicron and ancestral Wuhan variants on the rat model.

**Methods:** Viruses were isolated from a clinical specimen and virus characterisation performed. After inactivating the viral particles, the viruses were purified and formulated. Bivalent vaccines were a composition of 2.5 μg (5 μg total) or 5 μg (10 μg total) doses of each ansectral-based and Omicron-based monovalent vaccine. Subsequently, the potency of the monovalent and bivalent vaccines was investigated using the virus neutralisation test (VNT).

**Results:** The group that received three doses of the Omicron-specific vaccine demonstrated neutralisation activity against the Omicron variant with a geometric mean titer of 337.8. However, three doses of the Wuhan variant-specific vaccine could neutralise the Omicron variant at a maximum of 1/32 serum dilution. The neutralisation activity of the Omicron-specific vaccine, when administered as the booster dose after two doses of the Wuhan variant-specific vaccine, was 100% against the Omicron variant and the Wuhan variant at 1/64 and 1/128 serum dilution, respectively. Three doses of 5 μg bivalent vaccine could effectively neutralise both variants at the minimum of 1/128 serum dilution. The 10 μg bivalent vaccine at three doses showed even higher neutralisation titers: geometric mean titer of 338.0 against Omicron and 445.7 against Wuhan).

**Conclusion:** It is shown that the candidate bivalent vaccine could elicit a potent immune response against both Wuhan-Hu-1 and Omicron BA.1 variants. Therefore, we plan to evaluate the updated vaccine in the clinical trial setting.

## Introduction

The Omicron lineage of SARS-CoV-2, first described in November 2021, spread rapidly to become globally dominant and has split into many sublineages (1,2). With its high transmissibility and multiple novel spike (S) protein mutations, Omicron could escape from naturally acquired immunity while also challenging the effectiveness of current COVID-19 vaccines (2,3). According to the WHO weekly epidemiological update on COVID-19 released on 19 January 2023, globally, the Omicron variant of concern, accounted for 99.9% of reported SARS-Cov-2 viral sequences in the past 30 days (4).

The health impacts of future SARS-CoV-2 transmission depend on evolving mitigation strategies by health systems and societies. World Health Organization has reportedly declared that vaccination strategies based on repeated booster doses are unlikely to be sustainable. In this sense, the current COVID-19 vaccines originally developed against ancestral strains may need to be updated (5). The earliest attempts of boosting with Omicron-based vaccines had controversial results, some indicating similar efficacy to ancestral variant-based vaccine boosters in neutralising the Omicron variant (6), while others showed a 20-fold higher neutralising titers against the Omicron variant compared to wild-type boosters (7). Later findings indicated that bivalent vaccines containing both Omicron and ancestral strain-matched vaccines could elicit a superior neutralising antibody response against omicron and cross-reactivity with other variants, without evident safety concerns (8,9).

Since the beginning of the pandemic, Iran has endeavoured to develop and administer safe domestic vaccines, and considering the successful experiences of the country in the mass production of inactivated vaccines, applying this vaccine platform seemed the most feasible (10). BIV1-CovIran is an inactivated virus vaccine developed against the Wuhan variant of SARS-CoV-2. In phase I and II clinical trials, all adverse events were mild or moderate and transient. The seroconversion rate of neutralising-antibody was 82.8% and the vaccine neutralised SARS-CoV-2 at 1/64 serum titers among 82.5% of 280 participants who received two doses of 5 μg vaccine in a 28-day interval (10). In the phase III clinical trial, which is currently under peer review, a two-dose vaccine regimen was well tolerated, with no safety concerns, and conferred 70.5% and 83.1% efficacy against hospitalization and ICU admission, respectively. A historical cohort study also showed the vaccine has reduced hospitalization by 86.4% and deaths by 98.3% (11).

Exploiting our previous experience in constructing the Wuhan variant-based vaccine and commensurate with the global efforts in updating previous vaccines against the omicron variant, the inactivated virus vaccine platform was re-employed to develop an Omicron variant-based vaccine titled BIV1-CovIran Plus within a month of virus isolation (12). In the clinical trial study, A 5μg booster dose of the Omicron-based vaccine induced superior Omicron-specific total IgG antibody and more potent neutralisation titer compared to ancestral strain-based vaccines (neutralisation rate of 66% vs. 25% in 1/16 serum dilution).

Considering the enhanced immunogenicity and cross-reactivity shown in bivalent SARS-CoV-2 vaccines, a bivalent vaccine candidate was also developed containing an equal amount of inactivated virus particles of the ancestral SARS-CoV-2 (Wuhan-Hu-1) and the Omicron variant (BA.1) in two doses of 5 μg (2.5 μg each) and 10 μg (5 μg each). Here, we investigated and compared the neutralising antibody induction of the ancestral variant-based BIV1-CovIran vaccine, the Omicron variant-based BIV1-CovIran Plus vaccine, and the novel bivalent vaccine candidate, BBIV1-CovIran, against the Omicron and ancestral Wuhan variants on the rat model.

## Materials and methods

### Overview

This study was conducted following our previous study on the preclinical evaluation of the BIV1-CovIran vaccine in the mice model (13). Here, we conducted an animal study in the rat model to evaluate and compare the Omicron-specific vaccine (BIV1-CovIran Plus) and the bivalent vaccine (BBIV1-CovIran) with the ancestral variant-based vaccine (BIV1-CovIran) against Wuhan-Hu-1 and BA.1 SARS-CoV-2 variants. Since the construction method of Omicron-specific vaccine introduced here has not changed from the previously generated BIV1-CovIran vaccine (13). Briefly, the Omicron variant was isolated and an inactivated vaccine was generated against it. The 5-μg and 10-μg doses of bivalent vaccine consisted of 2.5 μg or 5 μg from each of the ancestral Wuhan-Hu-1 or Omicron inactivated vaccines was also generated. Subsequently, the safety and immunogenicity of the generated vaccines against the Wuhan and Omicron variant were evaluated and compared with the ancestral-based vaccine in animal models.

### Virus isolation and vaccines preparation

In the biosafety level□3 facility, the Omicron variant virus was isolated from a nasopharyngeal swab and subsequently infected mono-layer Vero cells (ATCC# CCL81). After incubation, virus growth was detected and confirmed by cytopathic effects (CPE), gene detection and electron microscopy. After passage, the viral RNAs were extracted and converted to cDNAs to be sequenced and characterized. Multiplication kinetics analysis of viral particles in Vero cells were demonstrated by TCID50, 72 hours post-infection.

The virus particles were inactivated by BPL (1:1400 dilution) at 2–8°C for 20–24 hours. Moreover, to eliminate the possibility of remaining live virus particles, three blind passages were done and no CPE was detected (14). Finally, the inactivated viruses were purified by ultracentrifugation and chromatography and then, formulated with Alhydrogel ^®^ adjuvant 2% (Brenntag Biosector).

In this study, two types of monovalent and bivalent vaccines were prepared. The monovalent vaccines were composed of a 5 μg dose of the Wuhan-specific or the Omicron-specific vaccines. On the other hand, bivalent vaccines were a composition of 2.5 μg or 5 μg doses of each vaccine.

### Animal Studies

Nine groups of five female, eight weeks old Wistar rat models provided by the Laboratory Animal Science Department, Pasteur Institute of Iran, Karaj, Iran were prepared. A total of 45 rats were divided into nine groups (each containing 5 rats). Some groups received two doses of vaccines on days 0 and 28, while others received three doses on days 0, 28, and 56. A detailed description of the study groups stratified by the vaccine they had been received is demonstrated in Table 1. All animal studies were conducted under the approval of animal ethics committee guidelines of the Ministry of Health and Medical Education (Tehran, Iran; ethical code: IR.ACECR.IBCRC.REC.1399.016).

**Table 1.** The vaccine type, dose numbers, and the days of injection among study groups

| Group | Vaccine | Number of doses | Days of injection |
| --- | --- | --- | --- |
| WW | BIV1-CovIran (monovalent ancestral-based) | 2 | 0-28 |
| **OO** | BIV1-CovIran Plus (monovalent Omicron-based) | 2 | 0-28 |
| **WWW** | BIV1-CovIran | 3 | 0-28-56 |
| **OOO** | BIV1-CovIran Plus | 3 | 0-28-56 |
| **WWO** | 2xBIV1-CovIran + 1xBIV1-CovIran Plus | 3 | 0-28-56 |
| **BB (2.5)** | BBIV1-CovIran (bivalent vaccine: 2.5µg +2.5µg) | 2 | 0-28 |
| **BB (5)** | BBIV1-CovIran (bivalent vaccine: 5µg +5µg) | 2 | 0-28 |
| **BBB (2.5)** | BBIV1-CovIran (2.5µg +2.5µg) | 3 | 0-28-56 |
| **BBB (5)** | BBIV1-CovIran (5µg +5µg) | 3 | 0-28-56 |

### Virus neutralisation test

The conventional virus neutralising test (cVNT) was conducted on sera acquired from rat groups (15–17). The heat-inactivated sera were mixed with Omicron and Wuhan variants of SARS-CoV-2 in the Dulbecco’s Modified Eagle Medium (DMEM). The mixtures were then inoculated onto Vero cells in 96□well plates and the supernatants (containing neutralising antibodies) were removed at one hour post infection. The infected Vero cells were kept in DMEM and incubated for 48 h at 37 °C in a 5% CO2 incubator. CPEs were recorded under microscopes 72 h post□infection and the neutralising antibody titers were measured as values of the highest dilution that inhibited CPE formation in each well.

## Results

### Viral particles successfully infected the Vero cells to prepare the vaccine seed

The Variant of Concern (VOC) Omicron GRA (B. 1.1.529+BA) SARS-CoV-2 strain was successfully isolated in Vero cells from the clinical specimen. The isolated strain had more than 99.9% identity to the original Omicron variant. Multiplication kinetics analysis of viral particles in Vero cells demonstrated the peak of multiplication with 10^6^ TCID_50_, 72 hours post-infection (Figure 1-A). After four passages, the vaccine seed was prepared, and the virus growth was recorded by the presence of CPE (Figures 1-B and C).

**Figure 1.**
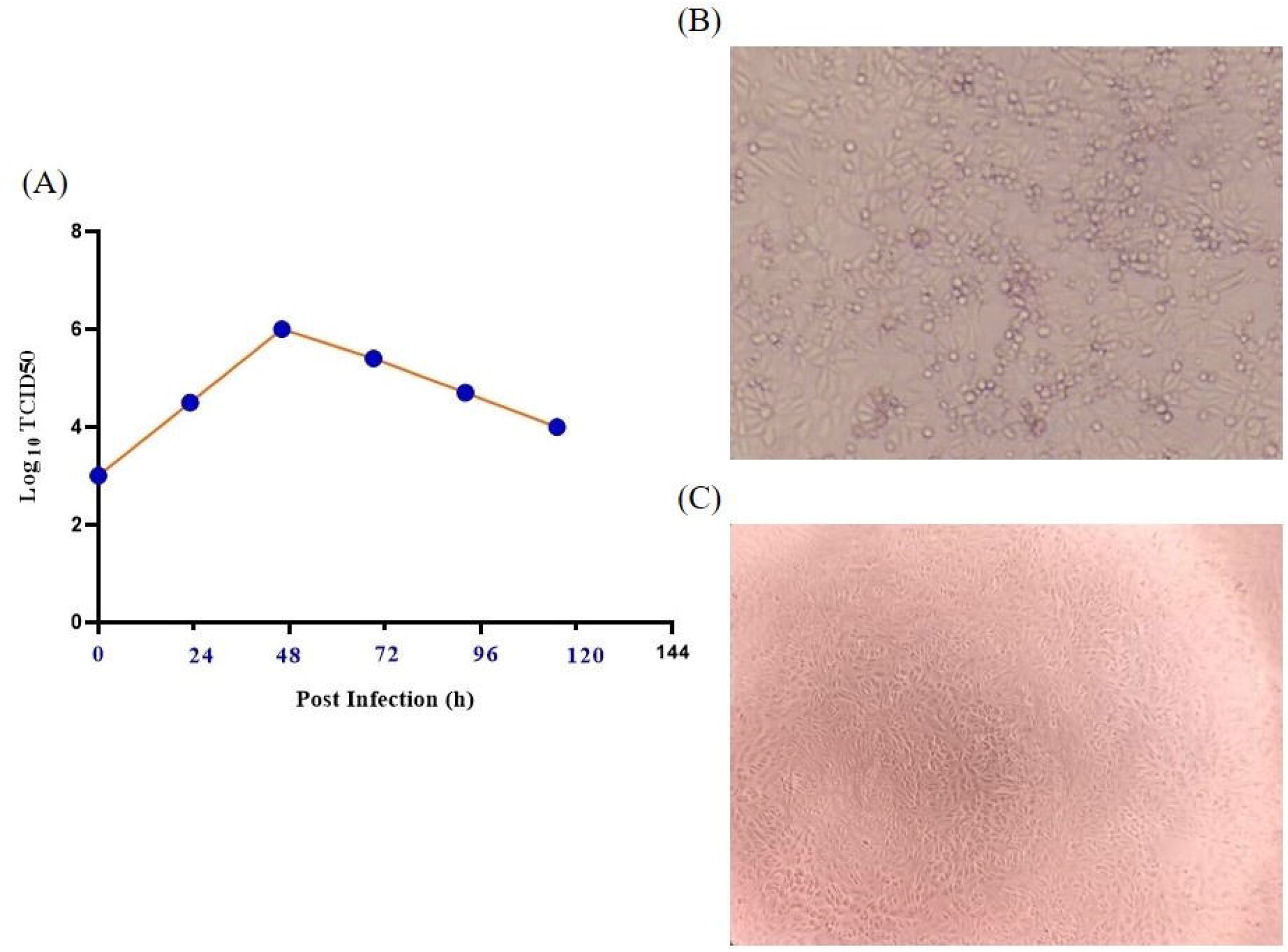
(A) The virus growth curve analysed by TCID50, (B) positive CPE resulting from virus infection, (C) CPE in the control sample.

### The monovalent Omicron-specific vaccine demonstrated neutralisation activity against the Omicron at various dose intervals

Among five rats that received two doses of the Omicron-specific on days 0 and 28, four of them (80%) demonstrated neutralisation activity against the Omicron variant at the 1/128 dilution (Figure 2A). On the other hand, two doses of Wuhan-specific could neutralise the Omicron variant at maximum 1/8 serum dilution (i.e., geometric mean: 97 vs. 7 against Omicron) (Figure 3). All of the samples’ sera (100%) in OO group neutralised the Wuhan variant at the 1/16 dilution (Figure 2B). In addition, among samples that received three doses of Omicron-specific on days 0, 28, and 56, the maximum dilution that successfully neutralised the Omicron variant was 1/512. Indeed, 3 out of five rats (60%) at 1/256 and 2 out of five (40%) at 1/512 sera dilution demonstrated neutralisation activity (Figure 2C). Three doses of Omicron-specific were also able to neutralise the Wuhan variant at 1/32 (80%) and 1/64 (20%) dilutions (Figure 2D). We also examined the neutralisation activity of the Omicron-specific vaccine when administered as the booster dose on day 56 after two doses of original the Wuhan-specific vaccine (i.e., BIV1-CovIran) on days 0 and 28. In this group, all five samples (100%) neutralised the Omicron variant and the Wuhan variant at 1/64 and 1/128 serum dilution, respectively (Figures 2C and D). The neutralisation potency in this group was also higher than three doses of Wuhan-specific vaccine against the Omicron variant (i.e., geometric mean: 64 vs. 27.9) (Figure 3).

**Figure 2.**
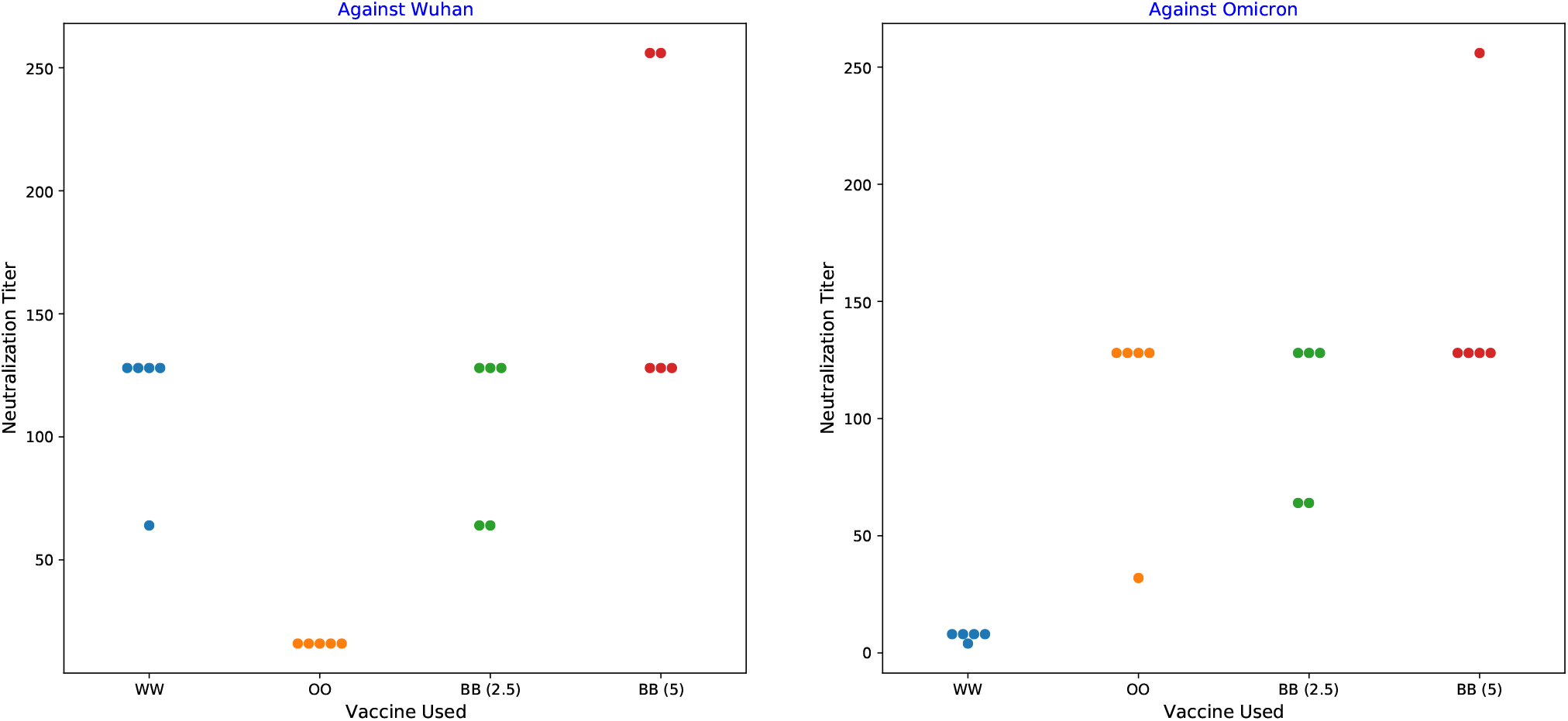
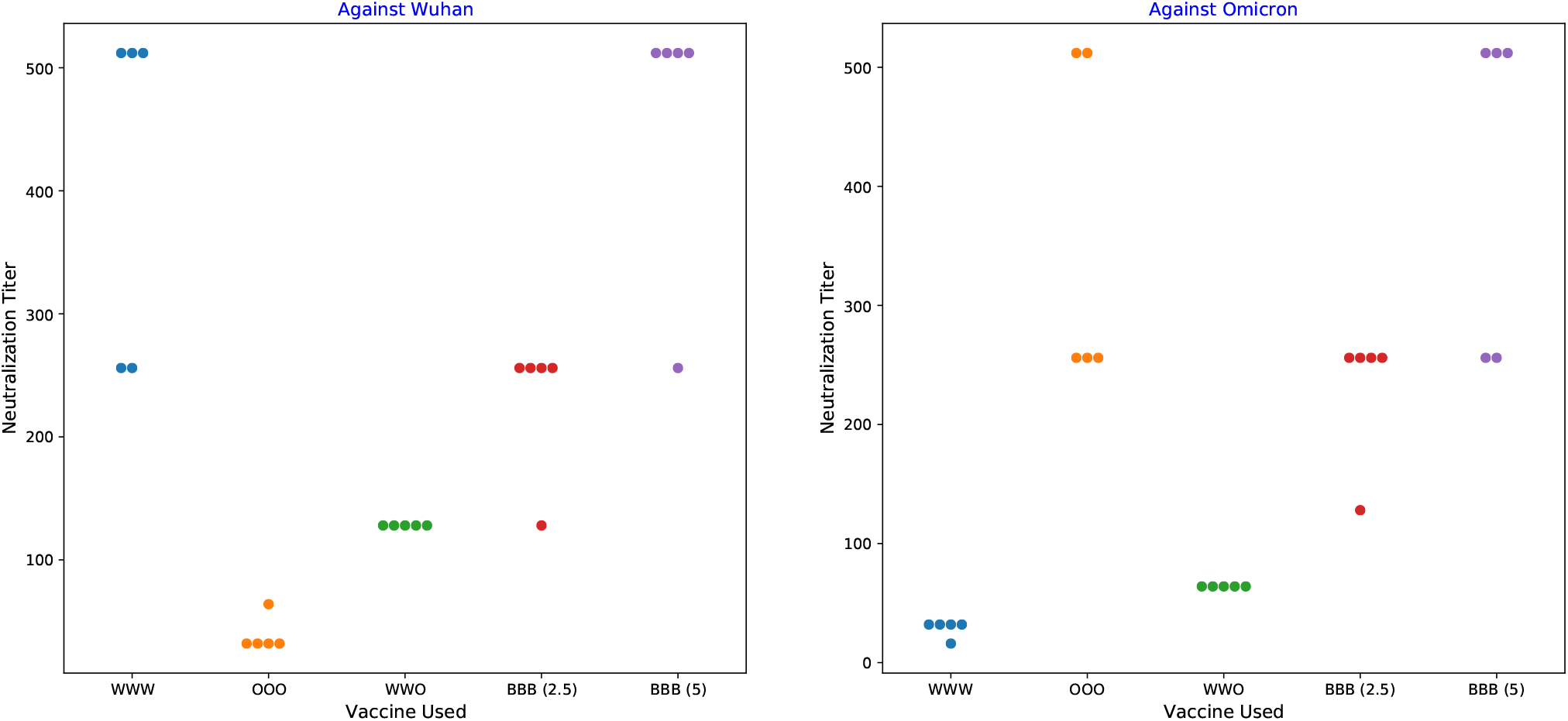
The neutralisation titer of samples against the Omicron and Wuhan variants (A) The neutralisation titer against the Wuhan variant in groups that received two doses of vaccines (B) The neutralisation titer against the Omicron variant in groups that received two doses of vaccines (C) The neutralisation titer against the Wuhan variant in groups that received three doses of vaccines (D) The neutralisation titer against the Omicron variant in groups that received three doses of vaccines WW: Two doses of BIV1-CovIran; OO: Two doses of BIV1-CovIran Plus; WWW: Three doses of BIV1-CovIran; OOO: Three doses of BIV1-CovIran Plus; WWO: BIV1-CovIran Plus booster after two doses of BIV1-CovIran; BB (2.5): Two doses of 5μg BBIV1-CovIran; BB (5): Two doses of 10μg BBIV1-CovIran; BBB (2.5): Three doses of 5μg BBIV1-CovIran; BBB (5): Three doses of 10μg BBIV1-CovIran

**Figure 3.**
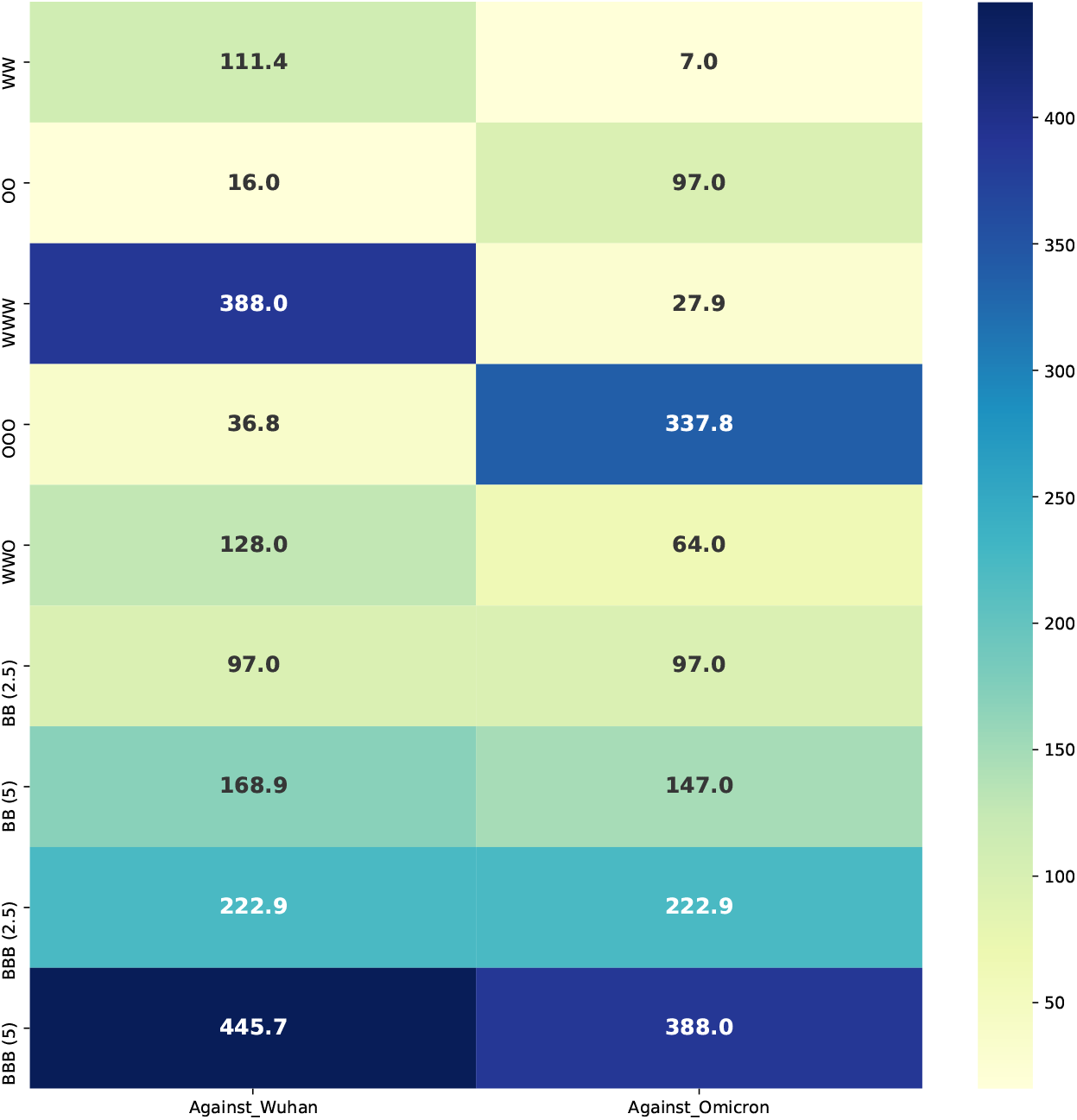
The geometric means of the neutralisation activity of study groups against the Omicron and Wuhan variants. WW: Two doses of BIV1-CovIran; OO: Two doses of BIV1-CovIran Plus; WWW: Three doses of BIV1-CovIran; OOO: Three doses of BIV1-CovIran Plus; WWO: BIV1-CovIran Plus booster after two doses of BIV1-CovIran; BB (2.5): Two doses of 5μg BBIV1-CovIran; BB (5): Two doses of 10μg BBIV1-CovIran; BBB (2.5): Three doses of 5μg BBIV1-CovIran; BBB (5): Three doses of 10μg BBIV1-CovIran

### The bivalent vaccine demonstrated neutralisation activity against both the Omicron and the Wuhan variants

Two different doses (i.e., 2.5 and 5 mg) of the Omicron-specific and the Wuhan-specific vaccines were mixed and injected into rats two or three times at 28-day intervals. The first group that had received two doses of the bivalent vaccine consisting of 2.5 mg from each vaccine on days 0 and 28 demonstrated similar neutralising activity against both variants (Figure 2). Accordingly, the sera of three rats (60%) could neutralise both the Omicron and Wuhan variants at 1/128 dilution, and the other two (40%) had neutralising activity at 1/64 serum dilution (i.e., geometric mean: 97) (Figure 3). When it came to the group that had received two doses of bivalent made from 5 mg of each vaccine, the minimum neutralising serum dilution was 1/128. 80% of the samples’ sera in this group neutralised the Omicron variant at 1/128 dilution, and 20% neutralised it at 1/256 (i.e., geometric mean: 147). On the other hand, 60% of them neutralised the Wuhan variant at 1/128 and 40% at 1/256 serum dilutions (i.e., geometric mean: 168.9). Finally, when we tried three doses of the bivalent vaccines on days 0, 28, and 56, the neutralising potency of sera was approximately one-fold increased. Four out of five rats (80%) that had received three doses of the 2.5mg mixture demonstrated neutralising activity against both the Omicron and the Wuhan variants at 1/256 serum dilution (Figure 4), and the other one at 1/512 dilution. In the last group, three doses of 5mg mixture were tested and the maximum neutralising serum dilution was 1/256. In this group, 60% and 80% of the samples’ sera could neutralise the Omicron and Wuhan variants, respectively at 1/512 dilution (i.e., the geometric mean of 445.7 against Wuhan and 388 against Omicron) (Figure 3).

## Discussion

Omicron is associated with a marked ability of reinfection or breakthrough infection even in populations with wide vaccine coverage, benefiting from its heavily altered antigenicity (1,18). Since the identification of the variant, evidence prompted the need for developing Omicron-specific vaccines (7). Recent Omicron sublineages, BA.4/5, are currently causing most cases of COVID-19 in many countries, and have been shown to resist even more than BA.1 and BA.2 against neutralisation by triple-dosed vaccinee serums (19). It is evident that protection against symptomatic BA.4/5 reinfection in those with a previous pre-omicron SARS-CoV-2 infection was 35.5%, while in those with a history of infection with a post-Omicron subvariant (including BA.1 or BA.2), the protection was 76.2% (20). This highlights the role of prior exposure to Omicron subvariants in having high protection against the BA.4/5 (21). The BQ and XBB subvariants, which are now rapidly expanding, are shown to further compromise the efficacy of current COVID-19 vaccines. The three shots of the ancestral-based vaccine showed far lower neutralization titers against BQ.1, BQ.1.1, XBB, and XBB.1 compared to the ancestral strain, with reductions of >37-fold to >71-fold. The variants also showed alarmingly high resistance to neutralization by serum of persons recently boosted with the new bivalent (covering BA. 5) mRNA vaccines (22). Nonetheless, CDC data showed bivalent booster doses provided 73% additional protection against COVID-19 hospitalization compared to the ancestral-based monovalent vaccine through September-November 2022 when BQ.1 was circulating in the US (23).

To date (as of January 21, 2023), more than 13.2 billion doses of vaccine have been delivered to people, and 5.52 billion people, i.e. 69.3% of the global population, received at least one dose of COVID-19 vaccine (Our World in Data; Coronavirus (COVID-19) Vaccinations). However, almost all those vaccines were designed based on the antigens from the ancestral virus. In this study, we showed that even three doses of the ancestral-based vaccine could neutralise Omicron with a geometric mean titer of 27.9; whereas, three doses of the Omicron-based vaccine can result in 12-fold higher titer. This was in line with our previous pilot animal study in the mice model to evaluate the neutralization potency of the Omicron-specific vaccine, BIV1-CovIran, against Omicron variant, which its outline are presented in supplementary material (Supplementary material 1). We also showed delivering the third dose of the Omicron-based booster instead of repeating an ancestral-based booster dose can result in a 2-folds higher neutralisation titer (geometric mean of 27.9 vs. 64). Considering that Omicron and its sublineages may become the persistent variants that humans might face even when the pandemic ends (24), this study overwhelmingly recommends including an omicron component in COVID-19 booster vaccines. Updating current vaccines would shape immune antigenic memory to have the lowest antigenic distance to the current circulating SARS-CoV-2 subvariants, and makes the immune system responses compatible with the circulating epitopes. The updated vaccines could also be applicable in immunologically naïve populations, such as children who reach the age of eligibility for getting COVID-19 vaccines (6).

On the other hand, viral evolution continues and escape variants emerge, vaccine neutralization potency against multiple variants is highly desirable (8). In this study, the bivalent vaccine, without sacrificing potency against the Wuhan variant, achieved significantly higher neutralising titers against Omicron than the ancestral-based vaccine. Two doses of 5 μg bivalent vaccine (2.5 μg each) neutralised both Omicron and Wuhan variants by the geometric mean titer of 97. Interestingly, the titer against Omicron equals the titer elicited by two doses of the Omicron-based vaccine and is 13-fold higher than the titer elicited by two doses of the Wuhan-based vaccine. Delivering the third dose of bivalent vaccine increased the titer by more than 2-folds for both Omicron and Wuhan variants. Moreover, delivering the 10 μg bivalent vaccine (5 μg each) resulted in even higher neutralisation titers against Omicron and Wuhan variants in both two-dose and three-dose regimens. The broad neutralising antibody response shown in this study is in accordance with findings from bivalent vaccine boosters of Moderna in preclinical studies (25). The bivalent Omicron-containing vaccine candidate of Moderna not only elicited higher spike-binding antibody responses against omicron BA.1 than mRNA-1273 (based on ancestral strain) but also showed a superior response against BA.4/5 subvariants and ancestral Wuhan-1 variant. Mechanisms underlying the increased antibody responses with bivalent vaccines have yet to be elucidated but may include generating new memory immune responses (26).

This study is subjected to some limitations. First, the study evaluated neutralization titers against the Wuhan-Hu-1 and BA.1 sublineage; but the neutralization potency of the vaccines against BA.2, BQ.1, and XBB.1 subvariants remained unexplored. However, the prospering preclinical and real-world results of the Wuhan variant-based (BIV1-CovIran) against pre-Omicron variants, combined with the high anti-Omicron neutralization titers elicited by the Omicron-based vaccine (BIV1-CovIran Plus), would be a promise that the bivalent candidate may enhance immunity against all currently circulating variants (11,13). As the second limitation, non-neutralizing antibodies and cross-reactive T-cell responses, both of which could influence protective immunity, were not taken into account in our analysis (27). Future experiments using other circulating and emerging strains and evaluating other components of immune system may be informative to determine the breadth of the vaccine protection.

Immunity induced by existing COVID-19 vaccines is subjected to waning over time (28,29), which in the case of the Omicron is particularly concerning since it is coupled with the variant’s ability to escape the humoral immune response elicited by ancestral variant-based vaccines (30,31). Altogether, the study iterate that the population’s waning immunity against SARS-CoV-2 should be recovered using Omicron-specific vaccines. This is especially important for Iran’s case, in which the new peak of COVID-19 caused by BA.2, BQ.1, and XBB was officially declared by the ministry of health on December 30, 2022 (32). Our findings indicate that Omicron-specific and bivalent vaccines may be a new tool in response to emerging variants, conserving neutralisation potency against Omicron sublineages. Accordingly, we are planning to evaluate the Omicron-based and bivalent vaccines according to the latest FDA guidance on the emergency use authorization for COVID-19 vaccines (33).

## Ethical approval

All animal models and maintenance procedures were under the approval of animal ethics committee guidelines of the Ministry of Health and Medical Education (Tehran, Iran; ethical code: IR.ACECR.IBCRC.REC.1399.016).

## Contributors

Conceptualization: A.A., Hamidreza J., M.T., M.LB., Hassan J.; Data curation: A.A., M.T.; Formal Analysis: A.A., M.T.,; Funding acquisition: A.A., M.T.; Investigation: Hamidreza J., Hassan J.; Methodology: A.A., M.T., Hassan J.; Project administration: Hassan J.; Resources: Hassan J., Hamidreza J.; Supervision: Hassan J., Hamidreza J.; Validation: Hassan J., Hamidreza J.; Writing – original draft: A.A.; Writing – review & editing: A.A., Hamidreza J., M.T., M.LB., Hassan J.

