## Supplementary Material 1 for "Omicron-Specific and Bivalent Omicron-Containing Vaccine Candidates Elicit Potent Virus Neutralisation in the Animal Model"

The VOC Omicron GRA (B.1.1.529+BA) SARS-CoV-2 strain first detected in Botswana/Hong Kong/South Africa was successfully isolated in Vero cells from the clinical specimen. Multiplication kinetics analysis of viral particles in Vero cells demonstrated the peak of multiplication with 10^6^ TCID_50_, 72 hours post-infection. After four passages, the vaccine seed was prepared, and the virus growth was recorded by the presence of CPE.

The study was carried out among female, six-to-eight-week-old BALB/c mice and Guinea pig animal models provided by the Laboratory Animal Science Department, Pasteur Institute of Iran, Karaj, Iran. A full human dose (0.5 mL) of the BIV1-CovIran Plus was injected intraperitoneally to five female mice and two Guinea pigs for abnormal toxicity reactions evaluation. The animals were observed for seven days for any evidence of ill-health. After seven days, a veterinary pathologist evaluated organs and tissues for any signs of adverse reactions (1).

For potency evaluation, four groups of ten mice received two doses of BIV1-CovIran Plus or placebo (phosphate-buffered saline) at 7-day and 14-day intervals. The conventional virus-neutralizing test (cVNT) was conducted on sera acquired from vaccinated mice groups seven days after the second injection. Virus-specific CPEs were recorded under microscopes 72h post‐infection, and the neutralizing antibody titres were determined as values of the highest dilution that inhibited CPE formation in each well. The presence of neutralizing antibodies was reported positive if the potency were ≥1:4 and reported protective if the potency were ≥ 1:16 (2–4). All animal models and maintenance procedures were under the approval of animal ethics committee guidelines of the Ministry of Health and Medical Education (Tehran, Iran; ethical code: IR.ACECR.IBCRC.REC.1399.016).

There was no evidence of abnormal clinical symptoms, altered water or food intake, weight loss, and macroscopic tissue alterations among the mice and Guinea pigs that received BIV1-CovIran Plus. Moreover, there were no microscopic changes in the animals' liver, kidney, heart, and lung samples.

The neutralization potency of two doses of BIV1-CovIran Plus was assessed against the Omicron variant. In all samples from the study group that received two doses of BIV1-CovIran Plus at a 7-day interval, the sera at ≥1/32 times dilution would neutralize the Omicron variant SARS-CoV-2. Similarly, the sera of all samples from the study group, which received two doses of BIV1-CovIran Plus at a 14-day interval, at ≥1/64 times dilution, would neutralize the Omicron variant SARS-CoV-2. Moreover, six out of ten (60%) of the samples in this group would neutralize the Omicron variant of SARS-CoV-2 at 1/128 times dilution. CPE formation was observed in all samples from the control group, and no neutralizing activity was detected at any sera dilutions (Figure 2). In conclusion, BIV1-CovIran Plus was well-tolerated in the animal models, and no safety concerns were raised. Moreover, the vaccine candidate elicited protective neutralization against the Omicron variant in 100% of mice studied a week after the second injection.

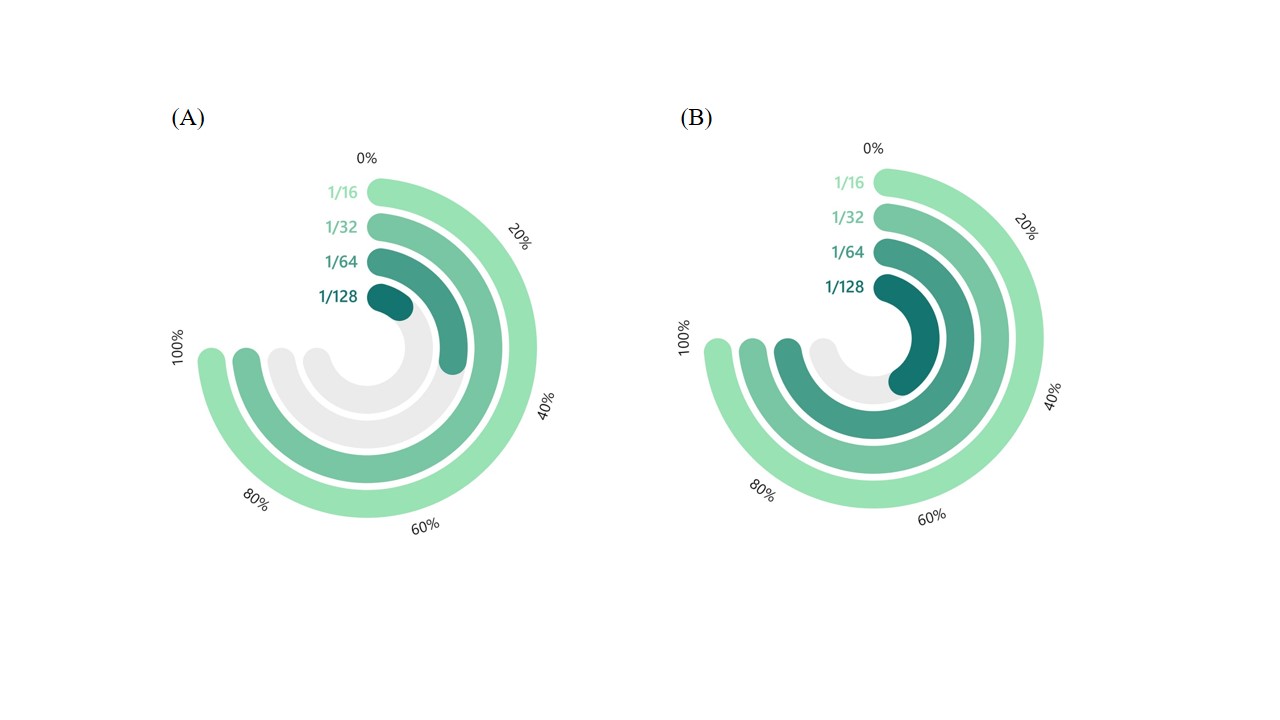
**Figure 2.** The virus neutralization titres seven days after receiving two doses of BIV1-CovIran Plus with 7 (A) and 14 (B) days intervals. 1/16 sera dilution was considered as the seroprotective threshold.
